# A comprehensive list of the replication promoters of *Bunyavirales* reveals a unique promoter structure in *Nairoviridae* differing from other virus families

**DOI:** 10.1101/2022.05.06.490981

**Authors:** Yutaro Neriya, Shohei Kojima, Arata Sakiyama, Mai Kishimoto, Takao Iketani, Tadashi Watanabe, Yuichi Abe, Hiroshi Shimoda, Keisuke Nakagawa, Takaaki Koma, Yusuke Matsumoto

## Abstract

Bunyaviruses belong to the order *Bunyavirales*, the largest group of RNA viruses. They infect a wide variety of host species around the world, including plants, animals and humans, and pose a major threat to public health. Major families in the order *Bunyavirales* have tri-segmented negative-sense RNA genomes, the 5’ and 3’ ends of which form complementary strands that serve as a replication promoter. Elucidation of the mechanisms by which viral RNA-dependent RNA polymerase recognizes the promoter to initiates RNA synthesis is important for understanding viral replication and pathogenesis, and for developing antivirals. A list of replication promoter configuration patterns may provide details on the differences in the replication mechanisms among bunyaviruses. Here, by using public sequence data of all known bunyavirus species, we constructed a comprehensive list of the replication promoters comprising 40 nucleotides in both the 5’ and 3’ ends of the genome that form a specific complementary strand. We showed that among tri-segmented bunyaviruses, viruses belonging to the family *Nairoviridae*, including the highly pathogenic Crimean-Congo hemorrhagic fever virus, have evolved a GC-rich promoter structure that differs from that of other bunyaviruses. The unique promoter structure might be related to the large genome size of the family *Nairoviridae* among bunyaviruses. It is possible that the large genome architecture confers a pathogenic advantage. The promoter list provided in this report is expected to be useful for predicting virus family-specific replication mechanisms of segmented negative-sense RNA viruses.

## Introduction

*Bunyavirales* is a new order that was recently proposed by the International Committee on Taxonomy of Viruses (ICTV). It consists of 12 families of closely related viruses: *Arenaviridae, Cruliviridae, Fimoviridae, Hantaviridae, Leishbuviridae, Mypoviridae, Nairoviridae, Peribunyaviridae, Phasmaviridae, Phenuiviridae, Tospoviridae*, and *Wupedeviridae*. Members of *Bunyavirales* have a segmented single-stranded negative-sense or ambisense RNA genome (1,2). The families *Arenaviridae, Hantaviridae, Nairoviridae, Peribunyaviridae*, and *Phenuiviridae* include several important pathogens that can cause severe diseases in animals, including humans, while the families *Fimoviridae, Phasmaviridae, Phenuiviridae*, and *Tospoviridae* include pathogens associated with plant diseases.

Major groups of bunyaviruses possess tri-segmented negative-sense RNA genomes, and share the same genetic organization consisting of three segments, *i.e*., the small (S), medium (M), and large (L) segments, based on their relative sizes. Each segment acts as a template for the replication of a positive-sense antigenome, and for the transcription of mRNA. The S segment encodes the nucleocapsid protein (NP), the M segment encodes a glycosylated polyprotein precursor (GPC) that is cleaved into envelope spike proteins Gn and Gc, and the L segment encodes the L protein, an RNA-dependent RNA polymerase (RdRp) responsible for the transcription and replication of the three RNA segments. The RNA synthesis activities of the three RNA segments are regulated by nucleotide (nt) sequences within the 3’ and 5’ untranslated regions (UTRs), which flank the S, M, and L open reading frames. The terminal nts of 3’ and 5’ UTRs exhibit complementarily, and such sequences have been shown to bind to, and influence the activity of, viral RdRp, promoting transcription to yield a 5’-capped mRNA by using cleaved host mRNA as a primer, and replication that results in the synthesis of a full-length copy of the genome template.

Bunyavirus promoters are composed of two promoter elements, *i.e*., promoter element 1 (PE1), the genomic extreme complement region, and PE2, the complement region located behind PE1, which was first described in Bunyamwera virus (BUNV) of the family *Peribunyaviridae* (3,4). PE1 comprises approximately 10 to 15 nts located at the extreme termini of the genome that are strictly conserved among all three segments. These nts have been shown to interact with L protein at separate sites in La Crosse virus (LACV) of the family *Peribunyaviridae* (5,6). PE2 comprises segment-specific nts at subsequent positions that are required to form canonical Watson-Crick base-pairing with corresponding nts at the opposite end of the template (3,4,7). These RdRp-RNA and RNA-RNA interactions are thought to account for the pseudocircular form of viral ribonucleoprotein complexes (8,9). In BUNV, sequence changes within PE1 have a significant effect on promoter function, but adjacent nts within PE2 are highly resistant to sequence changes, provided that their interterminal Watson-Crick base-pairing potential is maintained (3). For the family *Nairoviridae*, little is known on the roles of the 3’- and 5’-terminal UTRs in regulating RNA synthesis. As with other *Bunyavirales* members, the UTRs of all nairoviruses comprise highly conserved terminal proximal nts (PE1) shared by all three segments, followed by less conserved regions that are segment-specific nts (PE2). The importance of these segment-specific nts in RNA synthesis has been partially examined using a minigenome reporter assay in non-pathogenic *Hazara orthonairovirus* (HAZV), which is closely related to Crimean-Congo hemorrhagic fever virus (CCHFV) (10). PE1 and PE2 were found to be separated by a spacer region, which exhibited a critical requirement to be short in length and lack base-pairing ability. Taken together, the accumulated data indicate that the promoter structure of bunyaviruses differs among virus families. Understanding of these properties may be critical for developing antivirals targeting viral RNA synthesis and processing.

To characterize the promoter structure of diverse bunyaviruses, we constructed a list of all viral promoters that exhibit complementarity and each nt counts within the first 40 nts of both the 5’ and 3’ (anti)genomic ends. We found that the promoters of the family *Nairoviridae* differ from those of other virus families, and have characteristics unique to the family, which has a large genome size.

## Results

### Construction of the list of promoters in the order *Bunyavirales*

This study aimed to characterize the promoter structure of all virus species in the order *Bunyavirales*, including the families *Arenaviridae, Cruliviridae, Fimoviridae, Hantaviridae, Leishbuviridae, Mypoviridae, Nairoviridae, Peribunyaviridae, Phasmaviridae, Phenuiviridae, Tospoviridae*, and *Wupedeviridae*, which are tri-segmented or multi-segmented viruses, as summarized in Figure 1. The complementarity of the 5’ and 3’ extreme 40 nts of the genomic ends of virus genomes registered in the ICTV list was analyzed. Because there are incomplete genome sequences in the National Center for Biotechnology Information (NCBI) database that do not precisely cover the genome extremes, we selected viral sequences with complete complementarity in the terminal +1 to +3 nts (some exceptions with non-complementary +1 nt are included). The complement structure of 5’ and 3’ genomic ends (positive-sense form) as well as the genome length, counts of G:C/A:U complementarity, and counts of each nts (A, U, G and C) in the promoter region were calculated by using an automatic calculating system based on an Excel file (Supplementary Table 1), and the results are tabulated in Supplementary Table 2. A dataset was generated for each virus species that had complete data for all segments, including: tri-segmented bunyaviruses *Arenaviridae* (2 species), *Cruliviridae* (3 species), *Hantaviridae* (24 species), *Mypoviridae* (1 species), *Nairoviridae* (20 species), *Peribunyaviridae* (56 species), *Phasmaviridae* (2 species), *Phenuiviridae* (47 species), *Tospoviridae* (19 species) and *Wupedeviridae* (1 species); multi-segmented bunyaviruses *Fimoviridae* (17 species) and *Phenuiviridae* (14 species); and di-segmented bunyavirus *Arenaviridae* (23 species; characterized in Supplementary Figure 1).

**Figure 1.**
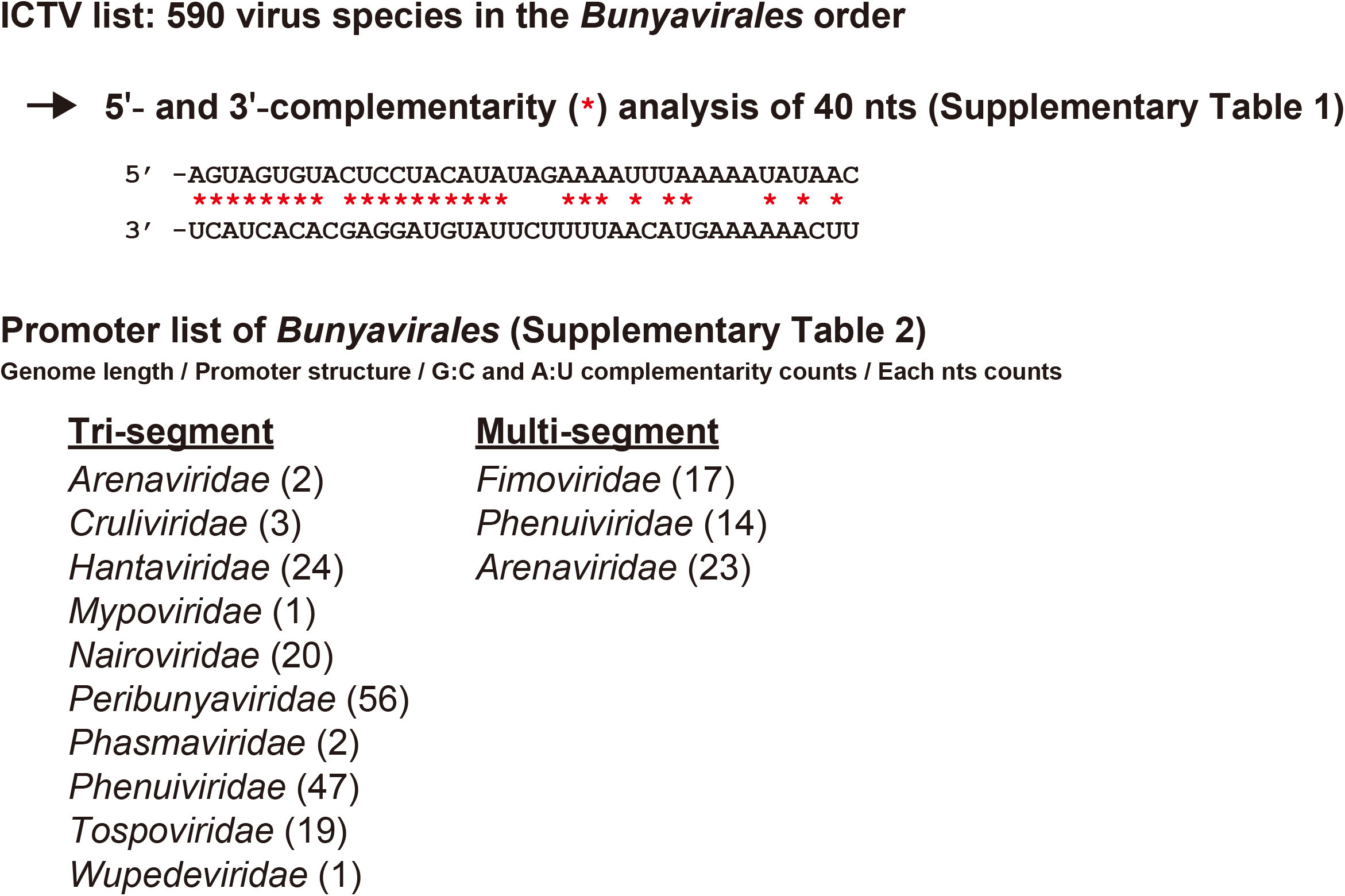
Construction of the promoter list of bunyaviruses. A schematic diagram of the promoter list construction. The promoter structure of 590 species of bunyaviruses registered in the ICTV list was analyzed. The genome length, promoter structure, G:C and A:U complementarity counts, and each nt (A, U, G, and C) count in the 40 nts of the promoters in the genomic ends are listed in Supplementary Table 2. The calculations were performed in an Excel file using automatic calculations for the promoter region as described in Supplementary Table 1.

### Characteristics of the replication promoters of five major virus families

For five major tri-segmented virus families, *i.e*., *Peribunyaviridae, Phenuiviridae, Tospoviridae, Hantaviridae*, and *Nairoviridae*, we examined the conservation of the nts in the 40 nts at the promoter region using the sequence generator <u>WebLogo</u> (Figure 2A, which shows representative M segments, and Supplementary Figures 2 and 3). The promoters differed among viruses: the initial nt was adenosine (A) in *Peribunyaviridae, Phenuiviridae* and *Tospoviridae*, and was uridine (U) in *Hantaviridae* and *Nairoviridae*. These promoters were further categorized into those starting with a tri-nt repeat (5’-AGUAGU and 5’-UAGUAG) and those starting with a di-nt repeat (5’-ACAC, 5’-AGAG and 5’-UCUC) (Figure 2A and B). We next examined the percentages of G:C and A:U complementarity at every nt position in the promoter region among the virus species in each virus family (Figure 2A). The complementarity conformation was remarkably different among virus families. G:C complementarity was relatively higher in the virus genomes with a promoter starting with U than in those with a promoter starting with A. The genomes of the families *Hantaviridae* and *Nairoviridae* contain high G:C complementarity at the 13-to 16-nt and 17-to 21-nt positions, respectively. In some viruses belonging to the family *Phenuiviridae*, a shift of 1 nt at the 10th position from the 5’ extreme appeared to increase the complementarity of the subsequent 5’ and 3’ ends (11), but it did not increase the total G:C complementarity frequency in the promoter region of *Phenuiviridae* (data not shown). It has been reported that HAZV has a promoter composed of two complementary regions of PE1 and PE2 separated by a spacer region formed by non-complementary sequences at the 13-to 16-nt position (10). We found the same feature in most virus species of the *Nairoviridae* family (Figure 2 and Supplementary Figure 3).

**Figure 2.**
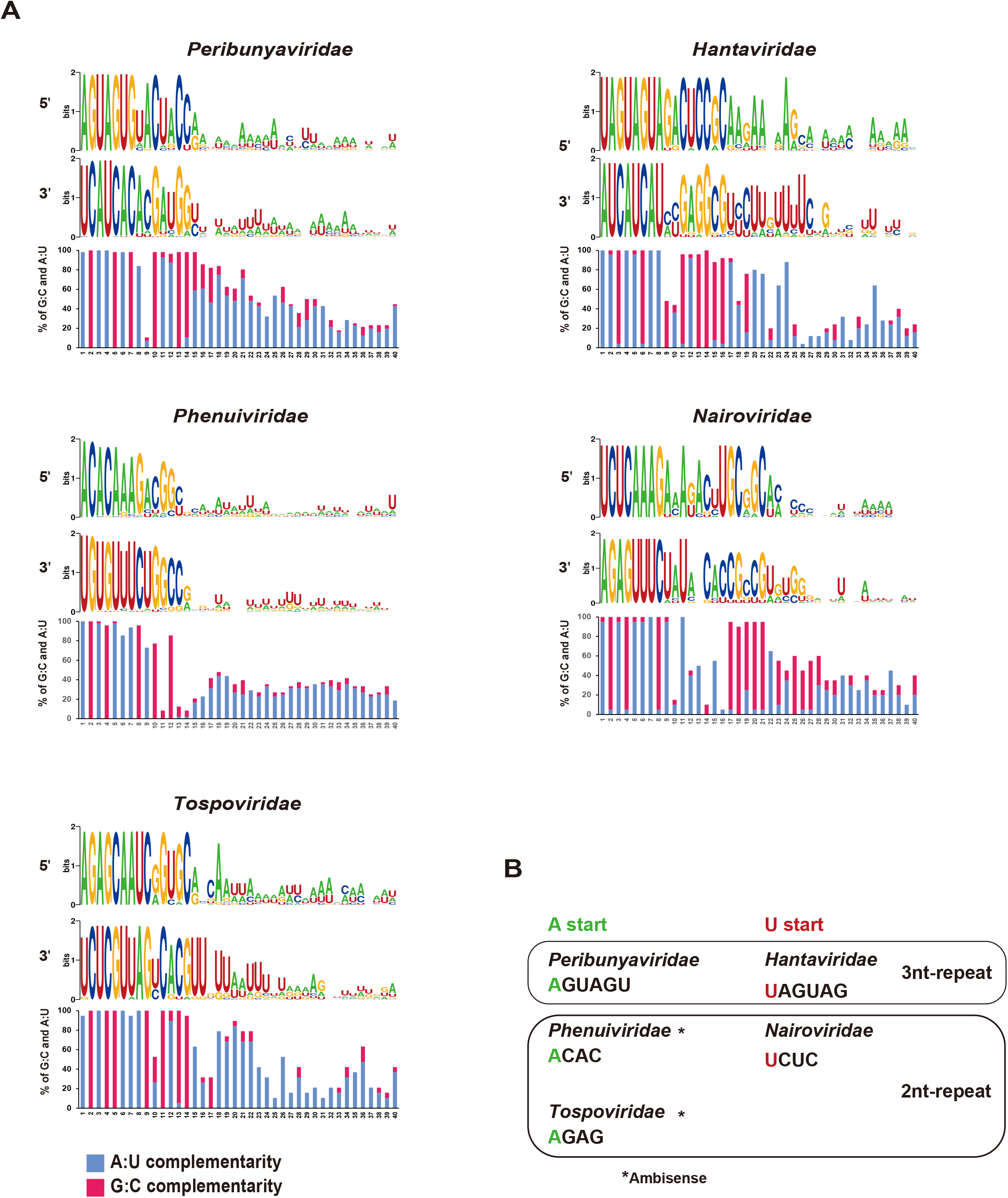
Characteristics of bunyavirus replication promoters in the M segment. (A) Sequence conservation and nt complementarity counts in the promoters of five major virus families. Sequence conservation was analyzed using the sequence logo generator WebLogo. The percentages of G:C and A:U complementarity in the promoter region (1 to 40 nts) among virus species in each family are shown as a bar graph. Virus species in the following families were examined: *Peribunyaviridae* (n = 56), *Phenuiviridae* (n = 47), *Tospoviridae* (n = 19), *Hantaviridae* (n = 24), and *Nairoviridae* (n = 20). (B) Characteristics of promoters. Virus promoters were divided into those that start with A and those that start with U, and further characterized according to the presence of a 3-nt or 2-nt repeat. *Phenuiviridae* and *Tospoviridae* possess an ambisense genome.

To analyze the promoter structure in more depth, the G:C and A:U complementarity in the 40-nt promoter region was determined in three segments of the five virus families. The average complementarity count of virus species in each virus family is shown in Figure 3A. The A:U complementarity counts were higher than the G:C complementarity counts in all segments for all virus families. However, G:C complementarity was particularly higher in the promoters of the family *Nairoviridae* than in those of other virus families (Figure 3A). Each nt (A, U, G, and C) in the promoter region was counted, and the average value within each virus family is shown in Figure 3B. In *Phenuiviridae, Tospoviridae*, and *Hantaviridae*, A in the 5’ end and U in the 3’ end were frequent in all segments. In *Peribunyaviridae*, both A and U were abundant at the 5’ and 3’ ends. In contrast, in the family *Nairoviridae*, C and G were more frequent at the 5’ and 3’ ends, respectively, than in other virus families. We demonstrated that the family *Nairoviridae* had more G:C complementarity as well as higher G/C counts in the 40 nts of the promoter region than other virus families, which is suggestive of stronger affinity for base pairing at both genomic ends.

**Figure 3.**
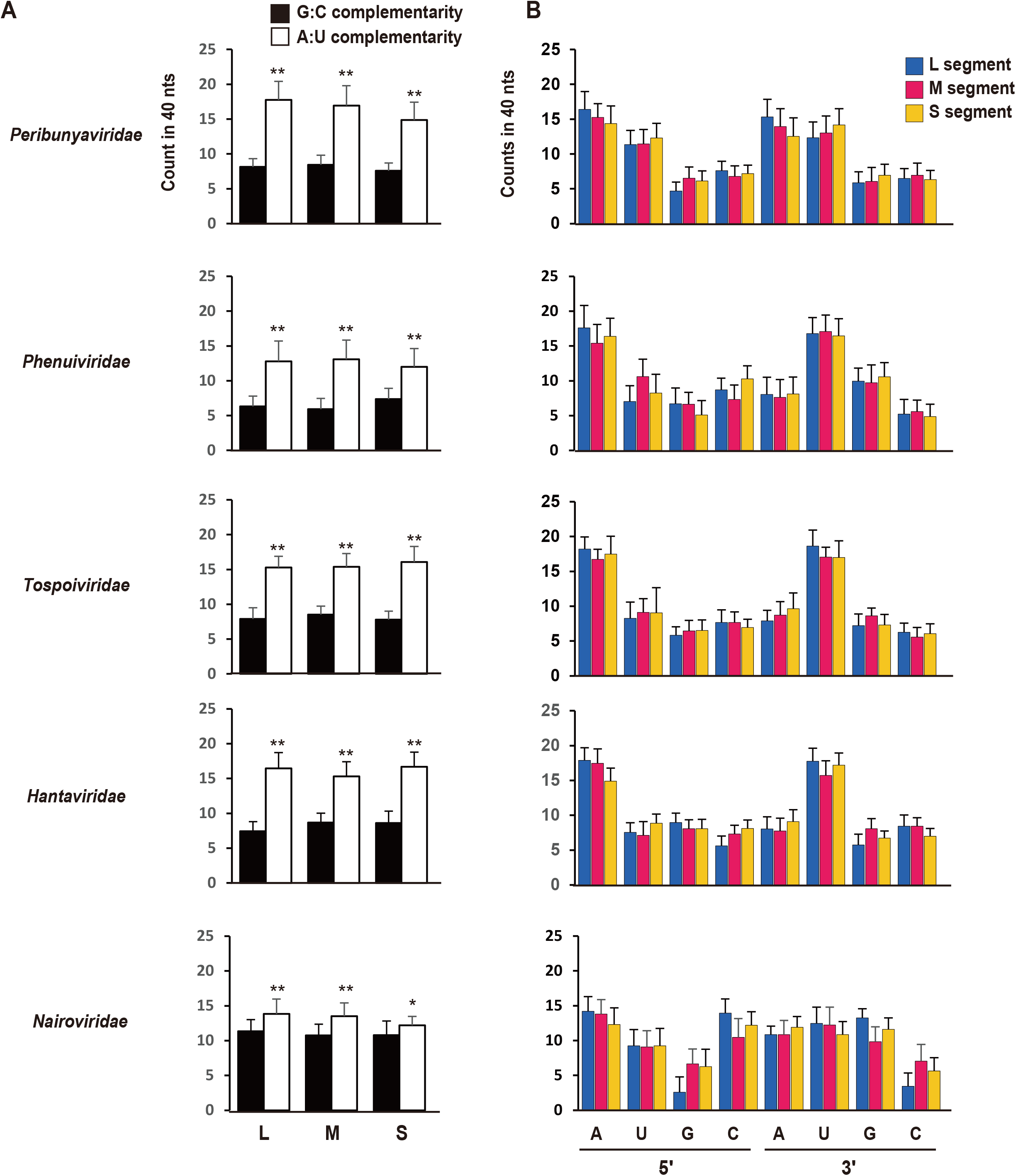
Counts of G:C and A:U complementarity, and each nt (A, U, G, and C) in the promoters. (A and B) The counts of G:C and A:U complementarity (A) and each nt (B) in the 40 nts of the promoters. Bars represent the means and standard deviations. Virus species in the following families were examined: *Peribunyaviridae* (n = 56), *Phenuiviridae* (n = 47), *Tospoviridae* (n = 19), *Hantaviridae* (n = 24), and *Nairoviridae* (n = 20). **p < 0.01, *p < 0.05, two-tailed unpaired Student’s *t*-test (G:C vs. A:U).

### Genome length of virus families in order *Bunyavirales*

The promoter structure of *Nairoviridae* differed from that of other tri-segmented virus families in that it had high G:C complementarity and a non-complementary nt spacer region. To investigate the relationship between these features and the characteristics of the viral genomes, the genome lengths of all viral species of the five virus families were studied. Figure 4A shows the average total genome length (combined length of the L, M, and S segments) of all virus species in the five virus families. The full genome lengths of families *Peribunyaviridae, Phenuiviridae*, and *Hantaviridae* were comparable, while the genome length of family *Tospoviridae* was larger, and that of family *Nairoviridae* was the largest. The length of each segment was also examined in all virus species, and the average lengths within virus families are shown in Figure 4B. Family *Tospoviridae* had relatively large L, M, and S segments. The L segment of *Nairoviridae* was the largest among all segments of all virus families.

**Figure 4.**
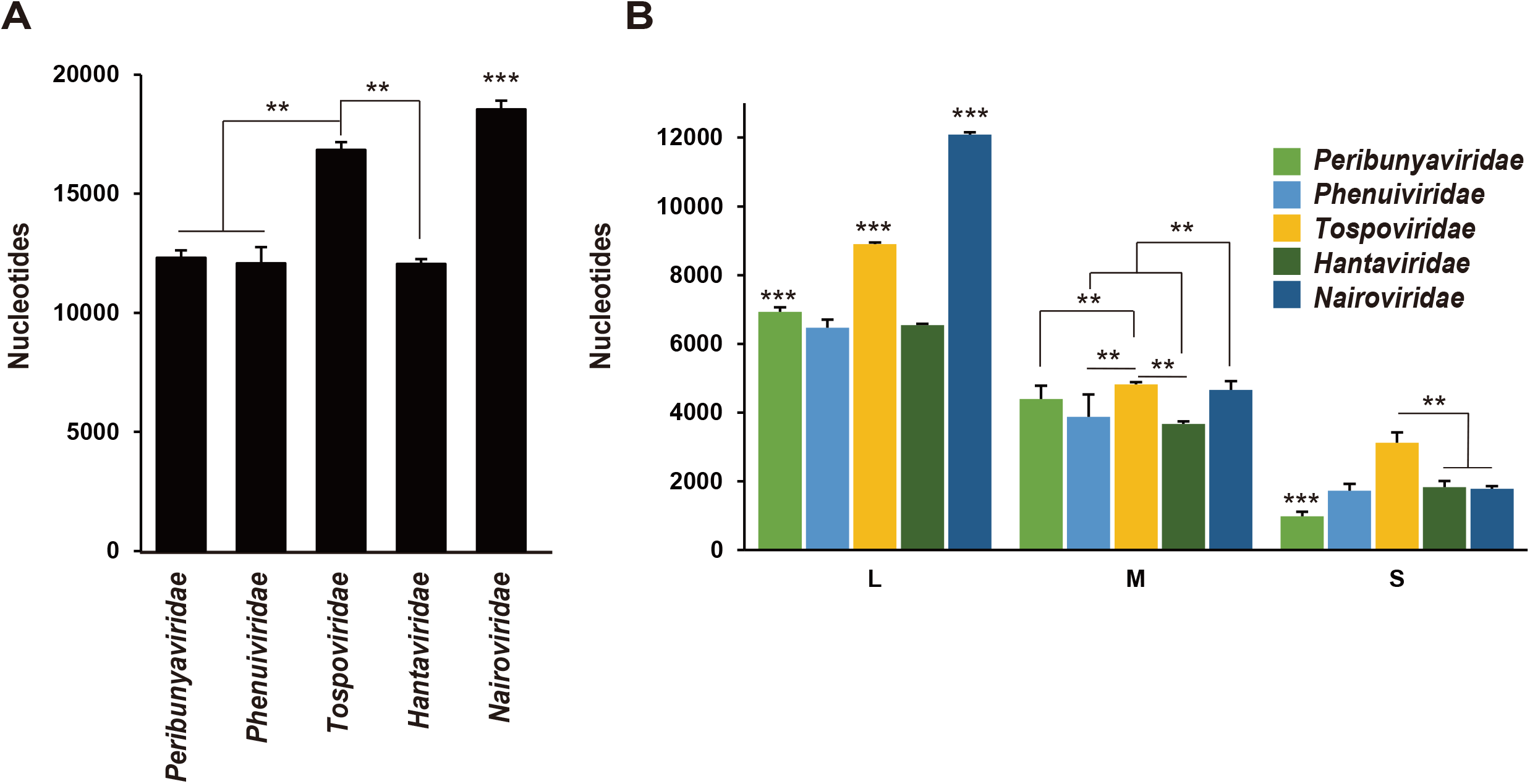
Genome length of bunyaviruses. (A) The total genome length (L segment + M segment + S segment) of each virus family. (B) The lengths of the L, M, and S segments in each virus family. Bars represent the means and standard deviations. Virus species in the following families were examined: *Peribunyaviridae* (n = 56), *Phenuiviridae* (n = 47), *Tospoviridae* (n = 19), *Hantaviridae* (n = 24), and *Nairoviridae* (n = 20). **p < 0.01, one-way ANOVA followed by Tukey’s test, *** p < 0.01, in comparison to the other four families.

### Genome length of virus species in family *Nairoviridae*

Among tri-segmented viruses belonging to the order *Bunyavirales*, only the family *Nairoviridae* includes highly pathogenic viruses categorized as biosafety level (BSL)-4 pathogens that cause hemorrhagic fever in humans, such as CCHFV (12). We hypothesized that the high pathogenicity of this virus family in mammals may be related to its large genome size. We examined the length of the available sequences annotated to “*Nairoviridae*” in the NCBI database. We first selected virus genome sequences possessing 5’-UCUC---GAGA-3’ ends, which are the most conserved genomic end sequences in nairoviruses, in the L, M and S segments. The lengths of these sequences are shown in Figure 5A. The family *Nairoviridae* contains two highly pathogenic viruses in mammals, *i.e*., CCHFV and Nairobi sheep disease virus (NSDV), which have a mortality rate of 30% and 90% in humans and small ruminants, respectively (13,14). The sequences of CCHFV and NSDV are shown in red and yellow bars in the graph, respectively (Figure 5A). The lengths of M segment of CCHFV and NSDV were all categorized in the largest group among virus genome sequences possessing 5’-UCUC---GAGA-3’ ends.

**Figure 5.**
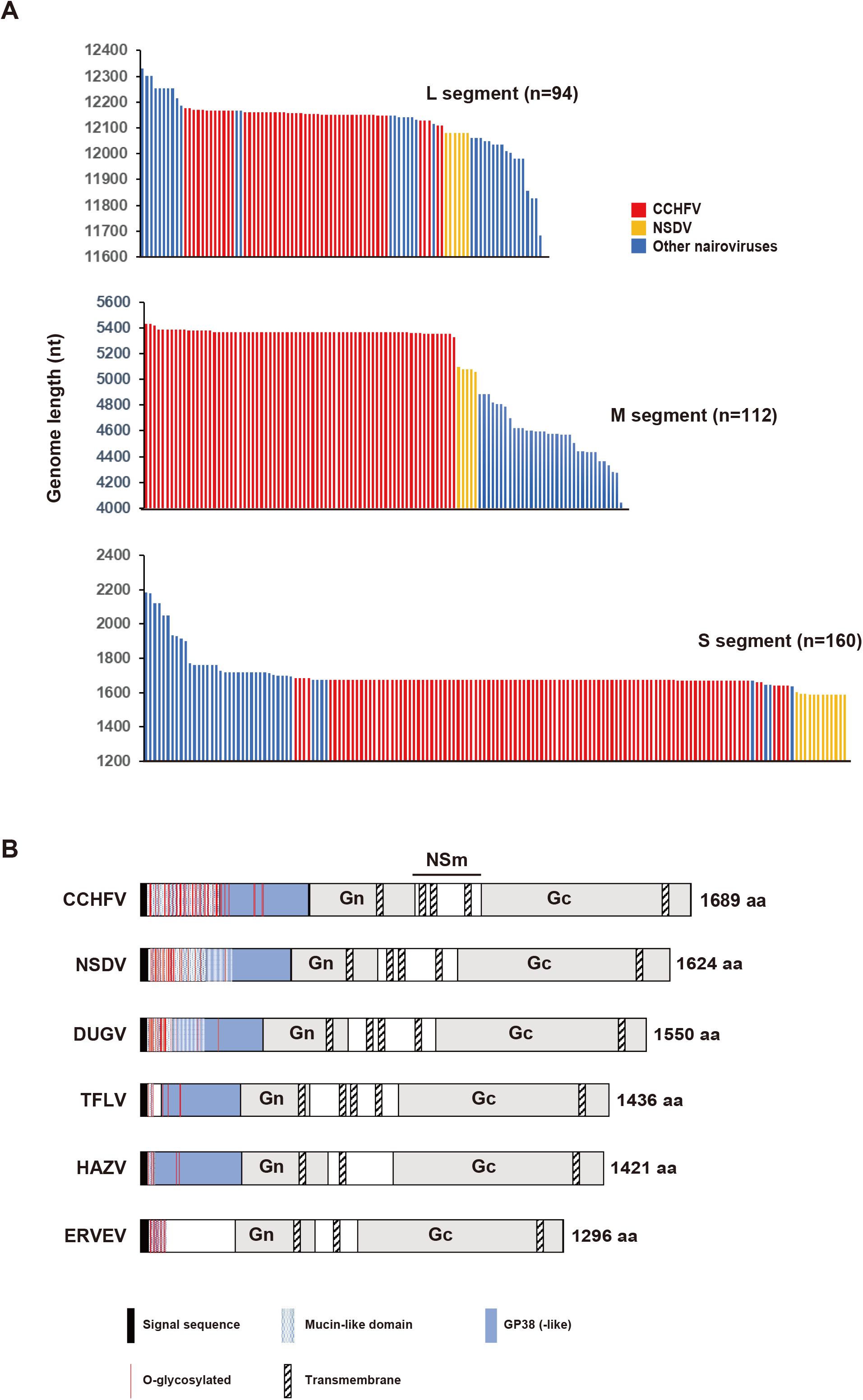
Genome length of virus species in the family *Nairoviridae*. (A) Genome length of each segment of nairovirus sequences (5’-UCUC---GAGA-3’) registered in the NCBI Refseq database. Bars represent the precise sequence length of each virus species. CCHFV and NSDV are shown in red and yellow. (B) Schematic diagram of GPC encoded in the M segment of nairoviruses. DUGV: Dugbe virus, TFLV: Tofla virus, ERVEV: Erve virus.

## Discussion

In this study, we tabulated the replication promoter structures of all known virus species in the order *Bunyavirales*. Our analysis focused on five major tri-segmented virus families, and the results indicated that the genomes can be divided into two categories: those with a promoter starting with A (families *Peribunyaviridae, Phenuiviridae*, and *Tospoviridae*) and those with a promoter starting with U (families *Hantaviridae* and *Nairoviridae*; Figure 2). Viral RNA polymerases have been shown to be able to initiate RNA synthesis with a purine (G or A), but not with a pyrimidine (U or C) (15). Therefore, the 5’-U genomic end of *Hantaviridae* and *Nairoviridae* is unconventional. The genomes of LACV and Rift Valley fever virus (RVFV) of the families *Peribunyaviridae* and *Phenuiviridae*, respectively, contain a 5’-triphosphate end that starts with A (5’-pppA) (16,17). The 5’-pppA is generated by viral RdRp that recognizes the opposite U as the template. As seen in several segmented and non-segmented RNA viral polymerases (18-20), bunyaviral RdRp synthesizes RNA from an internal nt, and not from the terminus of the template. In LACV, RNA synthesis is initiated with A using the U at the +4 position of the antigenome (3’-UCA<u>U</u>CA) as the template during genome replication (9). The elongated product, 5’-ppp<u>AGU,</u> is realigned to the +1 to +3 position of the antigenome template (3’-<u>UCA</u>UCA), and is further elongated to generate 5’-pppAGUAGU. Accordingly, the position of U responsible for RNA synthesis initiation is presumably the +3 position in the *Phenuiviridae* (3’-UG<u>U</u>G) and *Tospoviridae* (3’-UC<u>U</u>C) antigenomes. This indicates that the 5’-ppp<u>AC</u> and 5’-ppp<u>AG</u> products realign to the 3’-<u>UG</u>UG and 3’-<u>UC</u>UC of the antigenomes, respectively, and are further elongated to generate 5’-pppACAC and 5’-pppAGAG, respectively, which are precise complementary chains of the antigenome templates. In contrast, the genomes of Hantaan virus (HTNV) in the family *Hantaviridae*, and CCHFV in the family *Nairoviridae* contain a 5’-monophosphate end (16,21), suggesting an unconventional RNA processing event during replication. In HTNV, RNA synthesis is initiated with an internal G at the +3 position by using a C of the 3’-AU<u>C</u>AUC of the antigenome as the template (21). Subsequently, the elongated 5’-pppG<u>UA</u> product realigns to the 3’-<u>AU</u>CAUC to further produce 5’-pppGUAGUA. Then, the extreme 5’-pppG is removed by an endoribonuclease activity of viral RdRp to produce 5’-pUAGUA (5’-monophosphate end) (21). The endoribonuclease activity of RdRp is responsible for the cap snatching that cleaves the 5’ end of the host mRNA for use as a transcription primer (22). The −1 position of viral mRNA of HTNV is G, indicating that viral RdRp can cleave host mRNA after the G nt (cleave GpN to produce G/pN) during transcription. In *Nairoviridae*, the −1 position of viral mRNA is C (23,24), and it is also generated via the cap-snatching mechanism. Similar to the RNA synthesis in *Hantaviridae*, it is supposed that nairoviral RNA synthesis is internally initiated with 5’-pppC at the +2 position by using the G of 3’-A<u>G</u>A of the antigenome as the template. Subsequently, the 5’-pppC<u>U</u> product would realign to the 3’-<u>A</u>GA, and be further elongated to generate 5’-pppCUCU. The 5’-pppC would then be removed, resulting in the production of a 5’-monophosphate end. Therefore, although the hantaviral RdRp is a conventional enzyme that initiates RNA synthesis with a purine (G), the nairoviral RdRp is considered to be an unconventional enzyme that can initiate with a pyrimidine (C). Such a difference may be important for the targeting of novel antivirals specific for nairoviral diseases.

Our analysis additionally confirmed that most bunyaviral genomes begin with a di-or tri-nt repeat (Figure 2), which has been suggested previously (21,25). The repeats can determine the initiation site for RdRp (*e.g*., +2 in *Nairoviridae*, +3 in *Hantaviridae, Phenuiviridae* and *Tospoviridae*, and +4 in *Peribunyaviridae*), which is important for the prime-realign RNA synthesis mechanism. The biological significance of the internal position of RNA synthesis initiation is unclear. It is likely that the di-nt repeat is restricted to virus families possessing an ambisense genome, such as *Phenuiviridae* and *Tospoviridae*, as well as *Nairoviridae* (for which only CCHFV has been reported) (26). This suggests that the ambisense coding property may be related to the di-nt repetition in the genomic ends. If this is true, analysis of genomic end repetition patterns may enable the elucidation of new transcripts in various bunyaviral genomes.

The *Nairoviridae* promoter appears to have high G:C complementarity in the 17 to 21-nts region (Figure 2A and Supplementary Figure 2), and this likely reflects the high G:C complementarity rate at the promoter region (Figure 3A). Interestingly, this GC-rich dsRNA region is located after a spacer region composed of non-complementary bases around the 14th position in all three segments, as has been reported previously in HAZV and CCHFV (10,27). We have previously suggested the possibility that the HAZV polymerase can recognize this GC-rich dsRNA as a promoter element essential for RNA synthesis initiation via an unidentified domain of the L protein (10). This kind of specific protein-RNA interaction has been proposed to be a suitable target for antivirals against CCHFV, which is closely related to HAZV. Our comprehensive analysis of the promoter list also suggested that this kind of strategy may be applicable for all viruses belonging to the *Nairoviridae* family.

In bunyaviruses, genome replication in each segment is regulated by the segment-specific promoter strength, but the variations in nts (A, U, G, and C) in each promoter region do not differ significantly among the L, M, and S segments in all virus families, except for *Nairoviridae* (Figure 3B). It is possible that the promoter strength among segments is determined by slight differences in the promoter structure that do not affect the total complementarity counts or nt variations. Viruses in the family *Phenuiviridae* and CCHFV of the family *Nairoviridae* have an ambisense S segment, but there is no nt variation pattern in the promoter that is unique to the S segment (Figure 2B). This suggests that the nt variation in the promoter was not affected by the presence of the ambisense segment during the viral evolution process. On the other hand, the nt variation in the promoter of the nairoviral L segment was different from that of the M and S segments, *i.e*., it was observed to have less G and C at the 5’ and 3’ ends, respectively (Figure 2B). The nairoviral L segment is remarkably long when compared to other nairoviral segments and the genomes of other virus families (Figure 4B). This large genome size may be associated with the promoter structure.

It remains unclear why the genome of the family *Nairoviridae* is so large. *Nairoviridae* is the only tri-segmented virus family that includes hemorrhagic fever viruses classified as BSL-4 pathogens, such as CCHFV. We hypothesized that the large genome size of the family *Nairoviridae* may be related to its high pathogenesis in mammals. Although the length of the L segment in *Nairoviridae* is the longest among all bunyaviruses, it is not particularly long among the highly pathogenic viruses in this family (Figure 5A). Rather, our analysis confirmed that among viruses in the family *Nairoviridae*, the M segment is the largest segment in two highly pathogenic viruses in mammals, CCHFV and NSDV, suggesting that the M segment contains factors involved in viral pathogenesis. The M segment encodes GPC that is first translated as a polyprotein from mRNA, and further cleaved into Gn, Gc, and other accessory or uncharacterized proteins. A schematic diagram of several representative nairovirus GPCs is shown in Figure 5B. GPC contains an N-terminal signal peptide and multiple membrane-spanning domains, and is processed by signal peptidases to generate an N-terminal pre-Gn protein, C-terminal pre-Gc protein, and a double-membrane-spanning NSm protein. The pre-Gn and pre-Gc are subsequently processed by furin-like or subtilisin kexin isozyme-1 proteases to generate a mucin-like protein containing a large number of *O*-glycosylation sites, a protein designated as GP38 (-like), virion envelope glycoprotein Gn, and virion envelope glycoprotein Gc (28). We showed that although the sizes of Gn and Gc are similar among virus species, those of the *O*-glycosylation sites and GP38-like protein are different; in particular, they are larger in CCHFV and NSDV (Figure 5B). This suggests that these regions may be determinants of the pathogenicity of *Nairoviridae*. It has been proposed that GP38 is involved in CCHFV particle formation and viral infectivity (28). Analysis of convalescent patient sera showed high titers of CCHFV GP38 antibodies, which indicated the immunogenicity of this protein in humans during natural CCHFV infection. In a mouse model, an antibody against GP38 could protect the animals from a heterologous CCHFV challenge, indicating an association between GP38 and the high pathogenesis of CCHFV (29). Our present analysis indicates that there is an association between the N-terminal GPC region and viral pathogenesis not only in CCHFV, but also in other highly pathogenic nairoviruses, including NSDV.

In conclusion, we constructed a comprehensive list of the promoters in *Bunyavirales* that included all virus families in this order. Studies on the RNA synthesis mechanism of *Bunyavirales* have been limited to only a few virus species. Analysis of the conservation in all promoter structures is useful for the prediction of RNA synthesis mechanisms in uncharacterized and newly identified bunyaviruses. The automatic promoter-characterizing system (Supplementary Table 1) is applicable for all bunyaviruses for which the precise genomic end sequences are known.

## Methods

### List of bunyavirus promoters

In total, 590 bunyavirus species were registered in the ICTV list (https://talk.ictvonline.org/) on December 7th, 2021. The complete sequences of the L, M and S segments of bunyaviruses available on the NCBI associated with the GenBank accession numbers listed in Supplementary Table 2 were used for the analysis. After obtaining the full-length genome sequences, the sequences were input to the “Sequence” column in Supplementary Table 1, and the extreme 40 nts of each of the 5’ and 3’ ends, the complementarity between the 5’ and 3’ ends of the sequences, and the counts of G:C and A:U complementarity and each of the nts (A, U, G, and C) in the promoter region were calculated automatically. In Supplementary Table 1, the results of the L segments were input as representative. Conservation of the nts in the promoter was analyzed by using the sequence logo generator WebLogo (https://weblogo.berkeley.edu/logo.cgi).

### Analysis of the genome length of nairoviruses

Sequences annotated as “*Nairoviridae”* were downloaded from the NCBI refseq database on January 16th, 2022. There were 5,272 *Nairoviridae* sequences in the database. We first checked for the presence of the extreme promoter sequence 5’-UCUCA in the 8-nt ends of the sequences. The promoter sequence was present in both ends of 368 *Nairoviridae* sequences (Supplementary Table 3), and the lengths of these sequences were calculated using a custom Python script. The codes used for this analysis is available on GitHub (https://github.com/shohei-kojima/Arenaviridae_overhang_analysis_2022).

### Amino acid sequence map of the nairovirus glycoprotein

The structural characteristics of the nairovirus glycoprotein were predicted using TMHMM-2.0 for the transmembrane protein (30), SignalP-6.0 for the signal cleavage site (31), and NetOGlyc-4.0 for the O-linked glycosylation sites (32). Data on the glycoprotein sequences were collected from UniProt (https://www.ebi.ac.uk/uniprot/index). The UniProt accession numbers were: CCHFV, Q8JSZ3; NSDV, A0A0A7H8l1; Dugbe virus, Q02004; Tofla virus, A0A0U5AG15; HAZV, A6XIP3; and Erve virus, J3S7E1. GP38-like regions were found using the Protein Basic Local Alignment Search Tool (BLASTp) based on the amino acid sequences of the CCHFV and Dugbe virus GPC.

### Statistical analysis

Statistical analyses were performed with Prism software (version 9.1.2; GraphPad, San Diego, CA, USA). Statistical significance was assigned when p values were <0.05. Inferential statistical analysis was performed by a two-tailed unpaired Student’s *t*-test or one-way analysis of variance (ANOVA) followed by Tukey’s test, as appropriate.

## Author contributions

Y.N., S.K., A.S., M.K., T.I., T.W., Y.A., H.S., K.N., T.K. and Y.M. conceived and designed the study, performed the analyses, analyzed the data. Y.N. generated the automatic promotor calculator (Supplementary Table 1). S.K performed the genome length analysis of *Nairoviridae*. A.S. designed the schematic diagram of nairovirus GPC. Y.M. wrote the manuscript. All authors have read and agreed to the manuscript.

## Figure legends

**Supplementary Figure 1.**
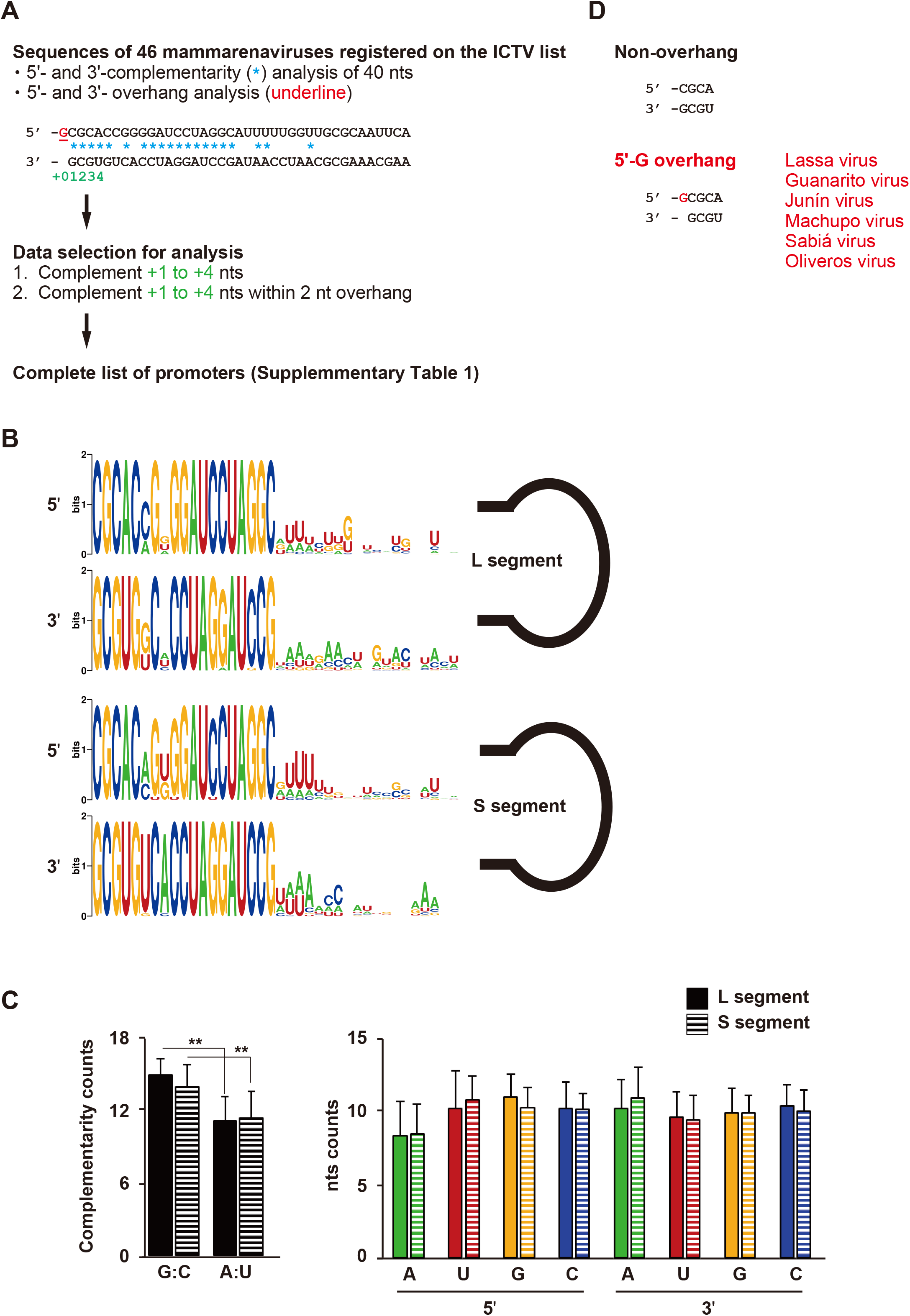
Construction of the list of promoters in *Arenaviridae*. (A) The genome of the family *Arenaviridae* possesses an unpaired nt at the genomic end that forms an overhang. To account for this overhang in the analysis, sequences that showed complementarity between +1 and +4 within 0-to 2-nt shifts in the 5’ or 3’ ends of all arenavirus genomes were selected. In total, 25 arenavirus promoter sequences, as listed in Supplementary Table 2, were included (2 tri-segmented antennaviruses, and 23 di-segmented mammarenaviruses). (B) The conservation of the extreme 38 nts (without the overhang) in each of the 5’ and 3’ ends of the L and S segments among the 23 mammarenavirus species was analyzed. (C) The counts of G:C and A:U complementarity, and A, U, G, and C in the first 40 nts (for 38 to 39 nts in the opposite strand of 2-nt and 1-nt overhangs, respectively) of the L and S segments were determined. G:C complementarity was significantly higher than A:U complementarity in both segments (**p < 0.01, one-way ANOVA followed by Tukey’s test), unlike in tri-segmented bunyaviruses. (D) A limited number of mammarenavirus species possessed an overhang nt in the database. It should be noted that many sequences of *Arenaviridae* genomes annotated in the NCBI database did not have overhanging genomic ends.

**Supplementary Figure 2.**
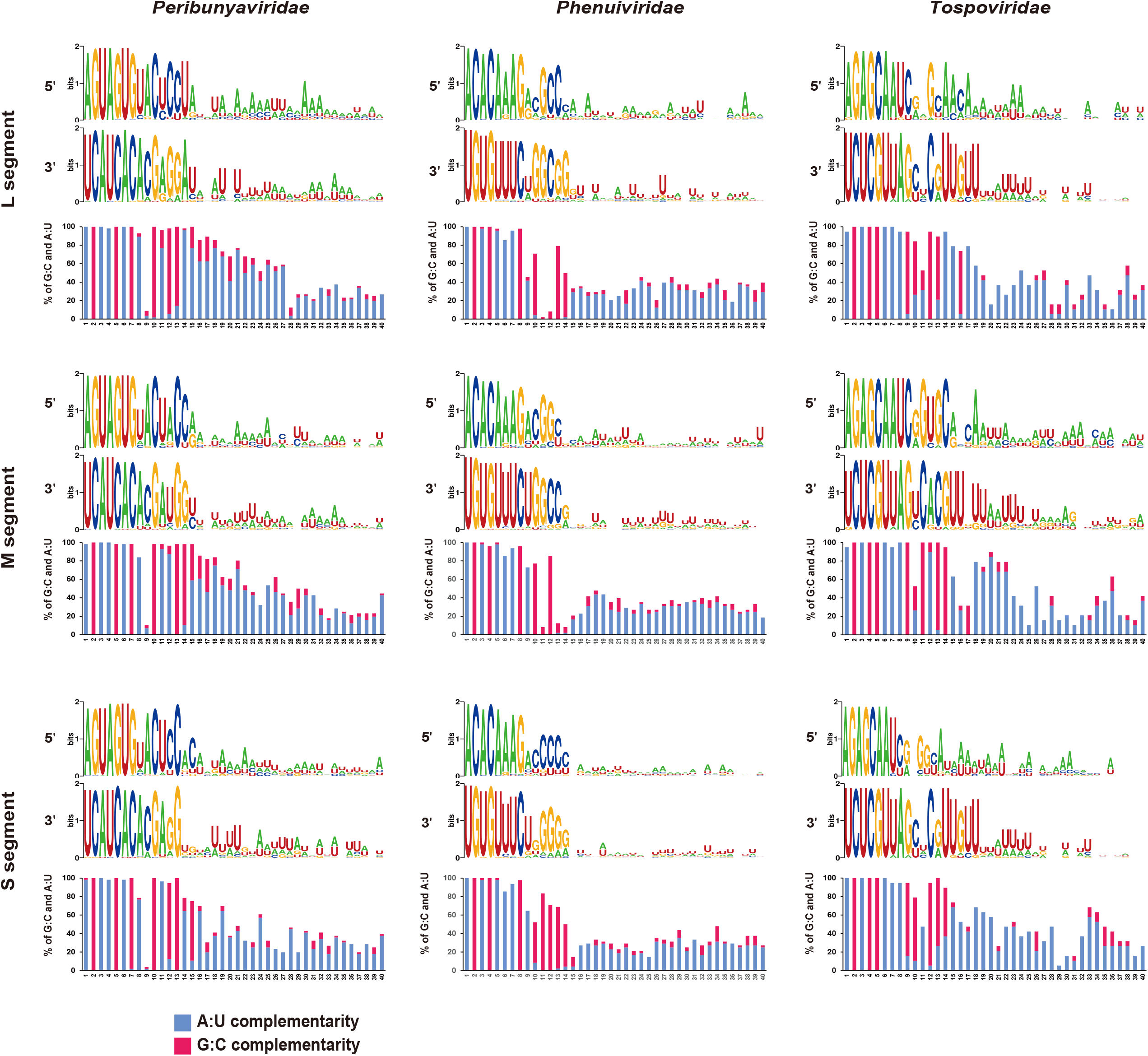
Characteristics of the replication promoters of the L, M, and S segments of *Peribunyaviridae, Phenuiviridae*, and *Tospoviridae*. Sequence conservation was analyzed using the sequence logo generator WebLogo. The percentages of G:C and A:U complementarity in the promoter region (1 to 40 nts) among virus species in each family are shown as a bar graph. Virus species in the following families were examined: *Peribunyaviridae* (n = 56), *Phenuiviridae* (n = 47), and *Tospoviridae* (n = 19).

**Supplementary Figure 3.**
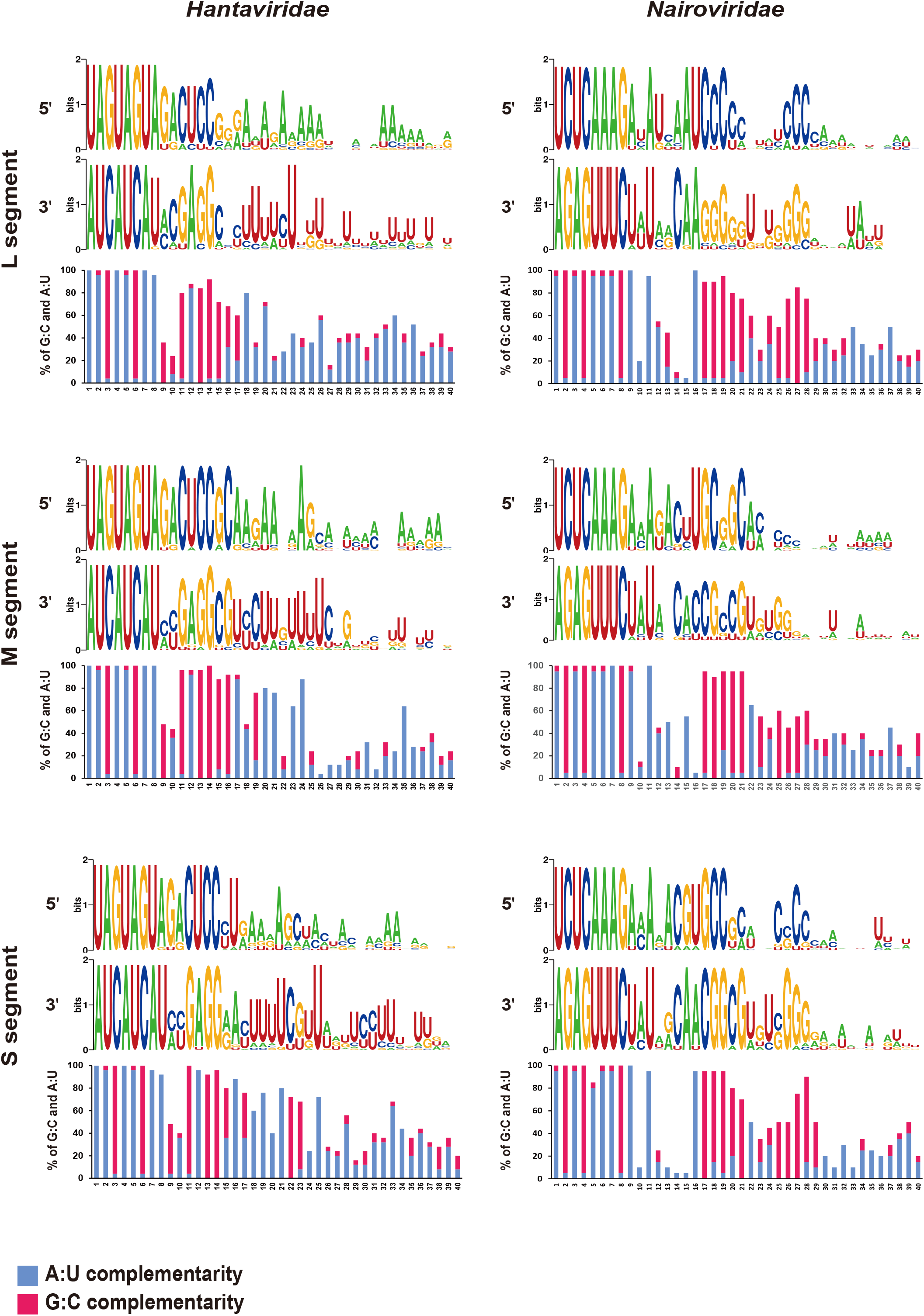
Characteristics of the replication promoters of the L, M, and S segments of *Hantaviridae* and *Nairoviridae*. Sequence conservation was analyzed using the sequence logo generator Weblogo. The percentages of G:C and A:U complementarity in the promoter region (1 to 40 nts) among virus species in each family are shown as a bar graph. Virus species in the following families were examined: *Hantaviridae* (n = 24) and *Nairoviridae* (n = 20).

**Supplementary Table 1. The automatic promotor calculator**

**Supplementary Table 2. Promoter list**

**Supplementary Table 3. *Nairoviridae* genome length**

